# Circular RNA Obelisk-*S.s* is highly abundant in *Streptococcus sanguinis* SK36

**DOI:** 10.1101/2024.03.24.586467

**Authors:** Rohan Maddamsetti, Lingchong You

**Affiliations:** Center for Quantitative Biodesign, Duke University, Durham, NC; Department of Biomedical Engineering, Duke University, Durham, NC

## Abstract

A new class of viroid-like circular RNAs, called Obelisks, was recently reported by Zheludev *et al*.^1^. They identify a specific 1137 nt Obelisk, called Obelisk-*S.s*, in monoculture transcriptomes of *Streptococcus sanguinis* SK36, a commensal bacterium of the human oral microbiome. Here, we report that Obelisk-*S.s*. is highly abundant in SK36, despite its absence from the SK36 genome (i.e., as DNA). In 11 out of 17 monoculture SK36 RNA-seq datasets examined, Obelisk-*S.s*. is more abundant than any mRNA. Given its abundance, we hypothesized that multiple Obelisk-*S.s* variants could coexist within SK36. We found three Obelisk-*S.s* mutations at 5-10% allele frequency in some samples: a R162R synonymous mutation in one set of replicate transcriptomes, and an I48I synonymous mutation and an intergenic mutation in another set of replicate transcriptomes. A simple mathematical model shows how high Obelisk abundance can transiently stabilize intracellular Obelisk populations, and how extreme Obelisk abundances may stabilize intracellular Obelisk populations indefinitely. Evolution experiments with SK36 could test this theory and could shed light on how Obelisks function and evolve within their microbial hosts.

## Introduction

In a recent preprint, Zheludev *et al*. report the discovery of a new kind of RNA, which they call Obelisks^1^. Obelisks are circular RNAs that are approximately 1000 nt in length, which fold into long hairpins and encode one or two proteins of unknown function. These unknown proteins are called Oblins. Obelisks are found globally and are prevalent in human oral microbiome metatranscriptomes^1^. Given the novelty of this discovery, we wanted to verify key aspects of this study.

In this report, we use independent transcriptomic data to confirm the existence of Obelisk-*S.s* in *Streptococcus sanguinis* SK36. Importantly, we found no evidence of Obelisk-*S.s* in SK36 genomic data, implying that Obelisk-*S.s* exists as an intracellular population of small circular RNAs within SK36 cells.

In addition, we report new aspects of Obelisk biology in SK36, namely its extreme abundance, and the presence of low frequency polymorphisms in Obelisk-*S.s* across RNA-seq technical replicates. These polymorphisms are not conserved across SK36 transcriptomes, suggesting that they arose independently during SK36 monoculture before cDNA preparation for RNA-seq. The evolutionary dynamics driving these preliminary patterns of Obelisk variation (i.e., mutation rates, strength of selection and distribution of fitness effects, Obelisk population bottlenecks during SK36 cell division and concomitant effects of genetic drift) remain a mystery in need of further investigation. Finally, we built a simple mathematical model to understand Obelisk persistence in SK36. The model shows that sufficiently high Obelisk abundance may stabilize Obelisk populations against loss, even in the absence of any fitness benefit to their host cell, given the assumption that Obelisks are randomly partitioned to daughter cells during cell division.

## Results

### *Obelisk-*S.s *is not found in the* S. sanguinis *SK36 genome*

We used NCBI BLAST to search for homology between Obelisk*-S.s* and the SK36 genome (Methods). No significant similarity was found. We repeated the search on the entire NCBI Nucleotide collection (nr/nt) database. Again, no significant similarity was found.

We asked whether this result was caused by errors in SK36 genome assembly. To do so, we downloaded Illumina resequencing data for SK36 from the NCBI Short Read Archive, and jointly mapped reads to the SK36 reference genome found in the NCBI RefSeq database (Accession: NC_009009.1) and the Obelisk-*S.s*. reference sequence (Methods). 100% of these sequencing reads mapped to the SK36 reference genome, with 225.7× sequencing coverage on average. By contrast, no sequencing reads at all mapped to the Obelisk-*S.s* reference sequence.

Together, these findings indicate that Obelisk-*S.s* is not found in the *S. sanguinis* SK36 genome.

### *Obelisk-*S.s *is highly abundant in the* S. sanguinis *SK36 transcriptome*

We then asked whether Obelisk-*S.s*. is found in the *S. sanguinis* SK36 transcriptome, as reported by Zheludev et al.^1^. We downloaded 17 RNA-seq runs collected from wildtype SK36 monoculture (Table 1). These data were collected and published by different research groups working independently^2-5^. Four of these datasets were analyzed by Zheludev et al. (2024). The remaining thirteen datasets were not; these independent data were submitted by researchers at Oregon Health & Science University in 2023 as part of two different projects^4,5^. I used *kallisto*^6^ (Methods) to map the Illumina reads from these RNA-seq datasets to the SK36 genome and the Obelisk-*S.s*. reference sequence (Supplementary Data File 1). Obelisk.*S.s*. is highly abundant in all of these samples (Table 1). Since RNA-seq determines relative and not absolute abundances for cellular RNA, I ranked RNA abundance for all 2270 protein-coding genes against Obelisk-*S.s*. in each sample. Ranking provides a robust and non-parametric measure of Obelisk-*S.s*. abundance in the SK36 transcriptome across independent samples. Obelisk-*S.s*. is more abundant than any mRNA in SK36 in 11 of the 17 RNA-seq datasets. Obelisk-*S.s*. is ranked no lower than 16 out of 2271 sequences in abundance in the remaining 6 RNA-seq datasets (Table 1).

**Table 1.** Wildtype *Streptococcus sanguinus* SK36 RNA-seq datasets analyzed for Obelisk-*S.s*.

| Bioproject<sup>1</sup> | SRA Run ID | Growth Condition<sup>2</sup> | RNA-seq Replicate<sup>3</sup> | Previously analyzed | RNA abundance rank of Obelisk-S.s.<sup>4</sup> |
| --- | --- | --- | --- | --- | --- |
| PRJNA270301 | SRR1713039 | BHI media, oxic | 1 | Yes | 1 of 2271 |
| PRJNA862079 | SRR20627698 | TY media + glucose, oxic | 1 | Yes | 10 of 2271 |
| PRJNA862079 | SRR20627697 | TY media + glucose, oxic | 2 | Yes | 6 of 2271 |
| PRJNA862079 | SRR20627696 | TY media + glucose, oxic | 3 | Yes | 6 of 2271 |
| PRJNA961761 | SRR24302371 | ASS media, oxic | 1 | No | 16 of 2271 |
| PRJNA961761 | SRR24302370 | ASS media, oxic | 2 | No | 13 of 2271 |
| PRJNA961761 | SRR24302369 | ASS media, oxic | 3 | No | 13 of 2271 |
| PRJNA937727 | SRR23591557 | CDM + sucrose, oxic | 1 | No | 1 of 2271 |
| PRJNA937727 | SRR23591555 | CDM + sucrose, oxic | 2 | No | 1 of 2271 |
| PRJNA937727 | SRR23591553 | CDM + sucrose, oxic | 3 | No | 1 of 2271 |
| PRJNA937727 | SRR23591551 | CDM + glucose, oxic | 1 | No | 1 of 2271 |
| PRJNA937727 | SRR23591549 | CDM + glucose, oxic | 2 | No | 1 of 2271 |
| PRJNA937727 | SRR23591547 | CDM + glucose, oxic | 3 | No | 1 of 2271 |
| PRJNA937727 | SRR23591545 | CDM + sucrose, anoxic | 2 | No | 1 of 2271 |
| PRJNA937727 | SRR23591543 | CDM + sucrose, anoxic | 3 | No | 1 of 2271 |
| PRJNA937727 | SRR23591541 | CDM + glucose, anoxic | 2 | No | 1 of 2271 |
| PRJNA937727 | SRR23591539 | CDM + glucose, anoxic | 3 | No | 1 of 2271 |

<sup>1</sup> PRJNA270301 was deposited in NCBI SRA in 2014 by Kyungpook National University.
PRJNA862079 was deposited in 2022 by the University of Florida. PRJNA961761 and PRJNA937727 was deposited in 2023 by Oregon Health & Science University.
<sup>2</sup> All samples were grown at 37C. BHI = Brain-Heart-Infusion, TY = Tryptone-Yeast, ASS = Artificial Saliva Solution, CDM = Chemically Defined Media. See references<sup>2-5</sup> for details.
<sup>3</sup> Replicate 1 for the CDM + sucrose, anoxic and CDM + glucose, anoxic conditions was not found in NCBI SRA, and therefore excluded.
<sup>4</sup> This comparison excludes rRNA and sRNA species; I only compare the abundance of reads mapping to the 2270 protein-coding genes in the SK36 genome (i.e., mRNA) to the abundance of reads mapping to Obelisk-S.s.

To check the robustness of this finding, I used *breseq* to map the Illumina reads from these RNA-seq datasets to the SK36 genome and the Obelisk-*S.s*. reference sequence (Table 2). Even though Obelisk-*S.s* (1137 nt) is less than 0.1% of the size of the SK36 genome (2,388,435 nt), at least 1% of all RNA-seq reads map to Obelisk-*S.s* in all samples.

**Table 2.**
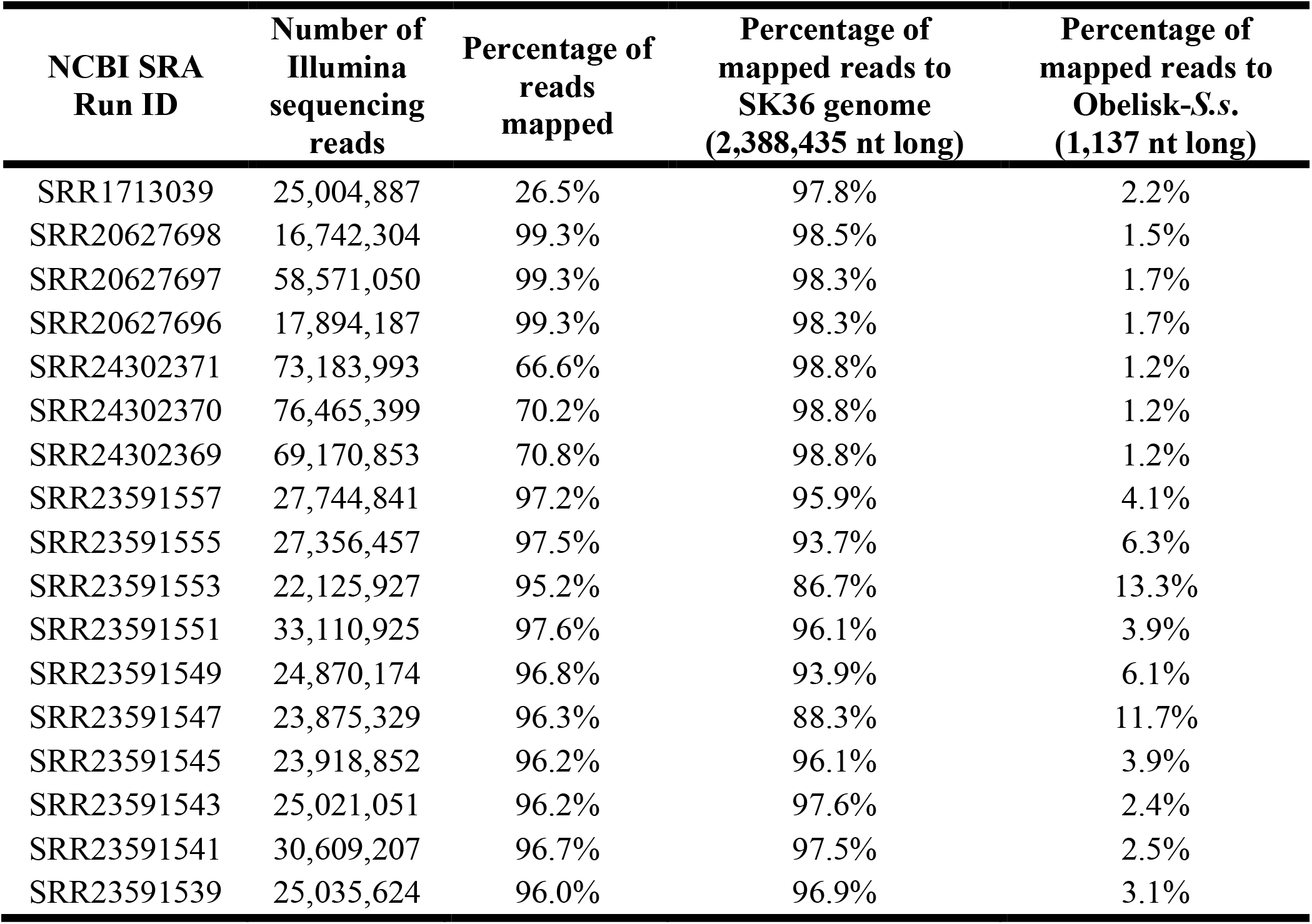
*breseq* RNA-seq read-mapping statistics.

| **NCBI SRA**<br>**Run ID** | **Number of**<br>**Illumina**<br>**sequencing**<br>**reads** | **Percentage of**<br>**reads**<br>**mapped** | **Percentage of**<br>**mapped reads to**<br>**SK36 genome**<br>**(2,388,435 nt long)** | **Percentage of**<br>**mapped reads to**<br>**Obelisk-S.s.**<br>**(1,137 nt long)** |
| --- | --- | --- | --- | --- |
| SRR1713039 | 25,004,887 | 26.5% | 97.8% | 2.2% |
| SRR20627698 | 16,742,304 | 99.3% | 98.5% | 1.5% |
| SRR20627697 | 58,571,050 | 99.3% | 98.3% | 1.7% |
| SRR20627696 | 17,894,187 | 99.3% | 98.3% | 1.7% |
| SRR24302371 | 73,183,993 | 66.6% | 98.8% | 1.2% |
| SRR24302370 | 76,465,399 | 70.2% | 98.8% | 1.2% |
| SRR24302369 | 69,170,853 | 70.8% | 98.8% | 1.2% |
| SRR23591557 | 27,744,841 | 97.2% | 95.9% | 4.1% |
| SRR23591555 | 27,356,457 | 97.5% | 93.7% | 6.3% |
| SRR23591553 | 22,125,927 | 95.2% | 86.7% | 13.3% |
| SRR23591551 | 33,110,925 | 97.6% | 96.1% | 3.9% |
| SRR23591549 | 24,870,174 | 96.8% | 93.9% | 6.1% |
| SRR23591547 | 23,875,329 | 96.3% | 88.3% | 11.7% |
| SRR23591545 | 23,918,852 | 96.2% | 96.1% | 3.9% |
| SRR23591543 | 25,021,051 | 96.2% | 97.6% | 2.4% |
| SRR23591541 | 30,609,207 | 96.7% | 97.5% | 2.5% |
| SRR23591539 | 25,035,624 | 96.0% | 96.9% | 3.1% |

Together, these analyses indicate that Obelisk-*S.s* is highly abundant in *S. sanguinis* SK36—even though it is not encoded as DNA in the SK36 genome.

### Evidence of Obelisk-S.s polymorphism

Although RNA-seq does not provide quantitative estimates of Obelisk*-S.s* copy number, it is evident that Obelisk*-S.s* exists at very high copy numbers in SK36 cells, relative to cellular mRNA transcripts. Given its high copy number, I asked whether single SK36 clones might contain diverse Obelisk subpopulations. By running *breseq* in polymorphism mode (Methods), I screened for segregrating mutations in Obelisk.*S.s*. To reduce false positives, I only considered mutations occurring above 5% allele frequency. The results are shown in Table 3. A R162R (CG<u>A</u>→CG<u>G</u>) *Oblin-1* mutation was found at 8-11% allele frequency in 2 out of 3 RNA-seq replicates in NCBI Bioproject PRJNA961761. A I48I (AT<u>C</u>→AT<u>A</u>) *Oblin-1* mutation and an intergenic (+274/–) mutation downstream of *Oblin-1* was found in several samples grown in different conditions in NCBI Bioproject PRJNA937727. By comparing Tables 1 and 3, one can see that these two mutations are always found in samples labeled as “Replicate 2” in this Bioproject. Therefore, it is likely that the apparent parallelism across replicates reflects some underlying correlation in how these samples were prepared, and not independent mutational events across independent SK36 cultures. For instance, it is possible that these Obelisk*-S.s* mutations evolved in the particular colonies or overnight cultures used to inoculate replicate cultures spanning different conditions for RNA isolation.

**Table 3.**
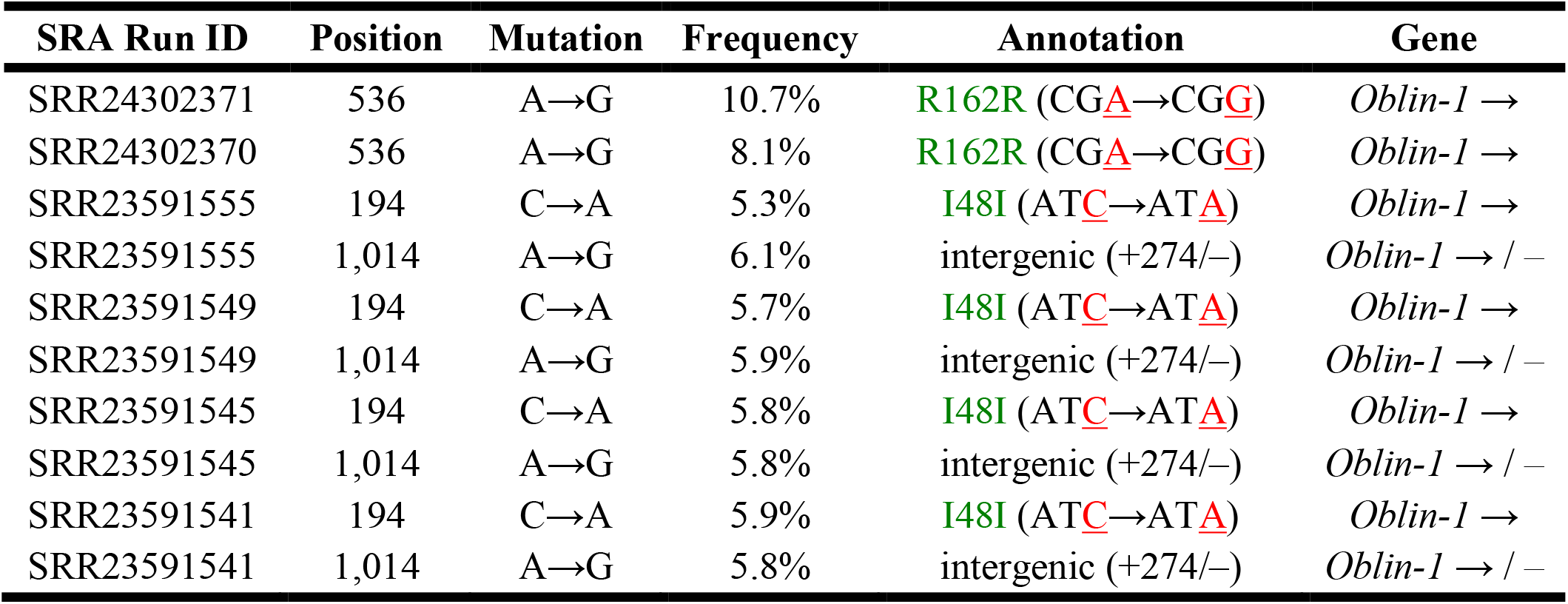
Obelisk-*S.s*. polymorphisms found across RNA-seq datasets.

| SRA Run ID | Position | Mutation | Frequency | Annotation | Gene |
| --- | --- | --- | --- | --- | --- |
| SRR24302371 | 536 | A→G | 10.7% | R162R (CG<u>A</u>→CG<u>G</u>) | *Oblin-1* → |
| SRR24302370 | 536 | A→G | 8.1% | R162R (CG<u>A</u>→CG<u>G</u>) | *Oblin-1* → |
| SRR23591555 | 194 | C→A | 5.3% | I48I (AT<u>C</u>→AT<u>A</u>) | *Oblin-1* → |
| SRR23591555 | 1,014 | A→G | 6.1% | intergenic (+274/-) | *Oblin-1* → / - |
| SRR23591549 | 194 | C→A | 5.7% | I48I (AT<u>C</u>→AT<u>A</u>) | *Oblin-1* → |
| SRR23591549 | 1,014 | A→G | 5.9% | intergenic (+274/-) | *Oblin-1* → / - |
| SRR23591545 | 194 | C→A | 5.8% | I48I (AT<u>C</u>→AT<u>A</u>) | *Oblin-1* → |
| SRR23591545 | 1,014 | A→G | 5.8% | intergenic (+274/-) | *Oblin-1* → / - |
| SRR23591541 | 194 | C→A | 5.9% | I48I (AT<u>C</u>→AT<u>A</u>) | *Oblin-1* → |
| SRR23591541 | 1,014 | A→G | 5.8% | intergenic (+274/-) | *Oblin-1* → / - |

### A mathematical model shows how high copy number can stabilize Obelisk persistence

The double-stranded hairpin secondary structure formed by Obelisk-*S.s* (Figure 1A) could help protect this RNA from cellular degradation^7^. It is easy to see that Obelisks could persist in SK36 if it confers some fitness benefit, or if its loss from the cell confers some fitness cost. In addition, its high cellular abundance could explain its persistence in SK36, even in the absence of any selective benefit. I examined the plausibility of this last hypothesis by building a simple mathematical model.

**Figure 1.**
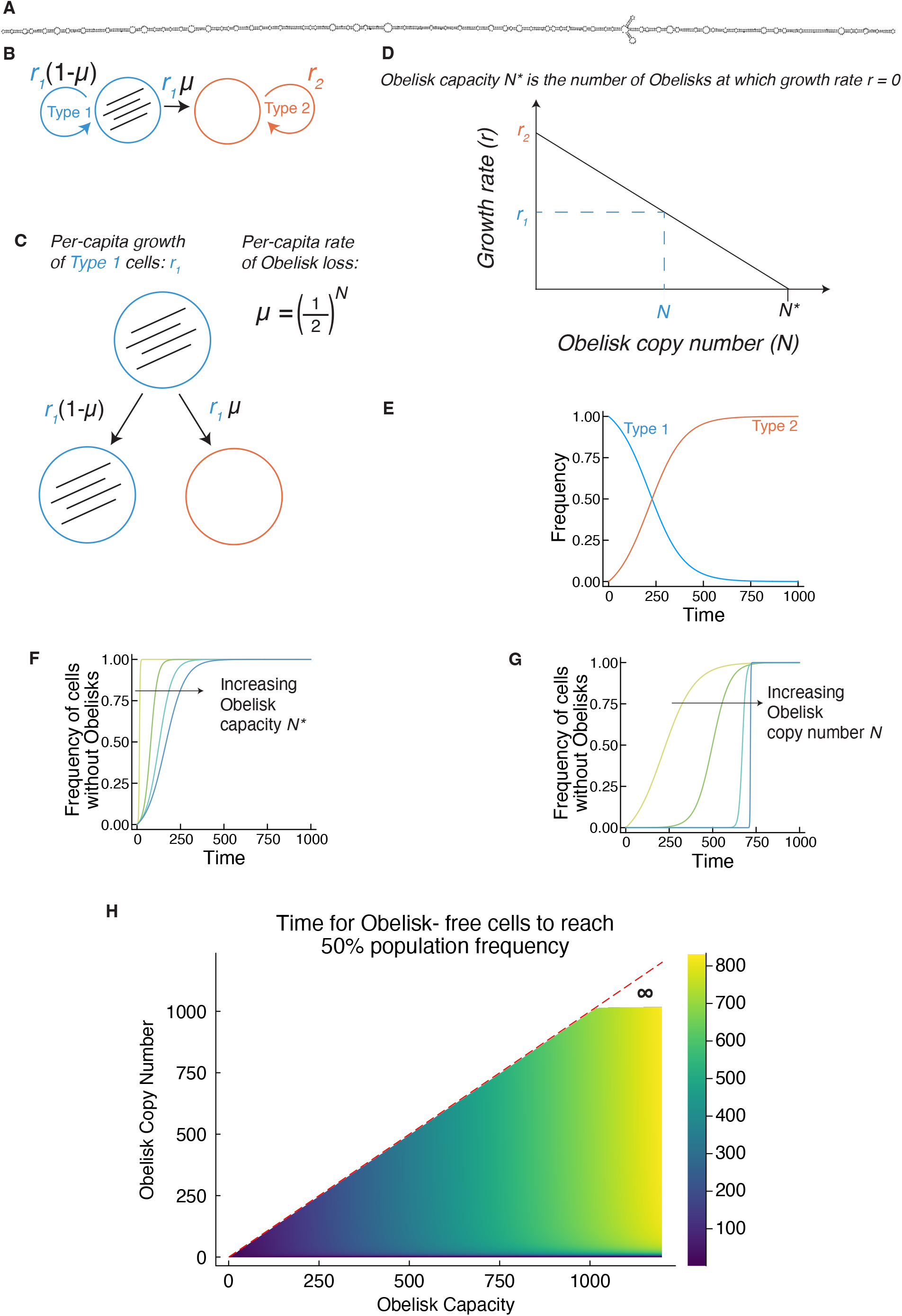
High Obelisk copy number can increase evolutionary stability by reducing the rate of generating Obelisk-free cells. A) RNAfold secondary structure of Obelisk-*S.s*. B) State diagram for the mathematical model. The two states represent cells containing intracellular Obelisks (Type 1), and cells that have lost intracellular Obelisks. C) Type 1 cells grow at rate *r*_*1*_. A Type 2 cell can be generated from a Type 1 cell at rate µ = (1/2)^N^, representing the probability of inheriting zero out of N Obelisks in its Type 1 parent due to random Obelisk segregation during cell division. Therefore, Type 2 cells are generated at rate *r*_*1*_µ. D) Obelisks are assumed to confer a small, linear fitness cost to the host cell. Obelisk capacity N* is defined as number of Obelisks in a cell at which growth rate *r*_*1*_ = 0. Increasing N* therefore reduces the per-capita fitness burden of an Obelisk. Type 2 cells have zero Obelisks and grow at rate *r*_*2*_. Type 1 cells have *N* obelisks and grow at rate *r*_*1*_. E) A typical simulation shows the evolution of Type 2 cells that outcompete Type 1 cells containing Obelisks. The simulation result in this panel uses the following parameter settings: *N* = 10, *N\** = 1000. F) Increasing Obelisk capacity *N\** decreases Obelisk fitness burden and has a small effect on the time for Type 2 cells to establish in the population. Colors shift from yellow to blue as *N\** increases. *N* is fixed at 10, and *N\** is varied from 20, 200, 400, 600. G) Increasing Obelisk copy number *N* increases Obelisk fitness burden but has a larger effect on the time for Type 2 cells to establish in the population. Colors shift from yellow to blue as *N* increases. *N\** is fixed at 1000, and *N* is varied from 10, 20, 100, 800. H) Increasing Obelisk copy number *N* and Obelisk capacity *N\** increase the evolutionary stability of Type 1 cells. When *N* is sufficiently high (∼1030 copies in numerical simulations), the per-capita Obelisk loss rate µ = (1/2)^N^ goes to zero, and Type 1 cells containing Obelisks become evolutionarily stable.

We model a population of cells containing Obelisks (Figure 1B). These cells grow at rate *r*_*1*_. A fraction µ of these newly produced cells lose the Obelisk due to random segregation at cell division. Therefore, µ = (1/2)^*N*^, where *N* is the Obelisk copy number (Figure 1C), and Obelisk-free mutants are produced at rate *r*_*1*_µ. Obelisks are assumed to confer a small, linear fitness cost to the host cell. We define Obelisk capacity *N*\* as number of Obelisks in a cell at which growth rate *r*_*1*_ = 0. Increasing N* therefore reduces the per-capita fitness burden of an Obelisk. Type 2 cells without obelisks grow at rate *r*_*2*_. Given particular values of the Obelisk copy number *N* and Obelisk capacity *N\**, it follows that *r*_*1*_ = *r*_*2*_ × [1 – (*N*/*N\**)] (Figure 1D).

These assumptions lead to the following linear system of ordinary differential equations (Methods):

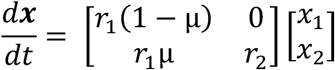

where **x** is a vector of two states, *x*_*1*_ (cells containing Obelisks) and *x*_*2*_ (cells without Obelisks). An example simulation of this model is shown in Figure 1E.

Increasing Obelisk capacity *N\** decreases Obelisk fitness burden and has a small effect on the time for cells without Obelisks to establish in the population, by reducing *r*_*1*_ (Figure 1F). Increasing Obelisk copy number *N* increases Obelisk fitness burden but has a larger effect on the time for cells without Obelisks to establish in the population, because the per-capita loss rate µ decreases geometrically with *N* (Figure 1G). By varying *N* and *N*\* and calculating the time for Obelisk-free cells to reach 50% population frequency (Methods), we find that when Obelisks reach biologically plausible copy numbers of ∼1200 Obelisks per cell, the per-capita Obelisk loss rate µ = (1/2)^N^ goes to zero, and Type 1 cells containing Obelisks become evolutionarily stable, despite the fitness cost conferred by abundant Obelisks.

## Discussion

Here, we confirm that Obelisk-*S.s*. indeed represents a novel class of viroid-like circular RNA in *S. sanguinis* SK36 and show that it is highly abundant in SK36 cells. As Obelisk-*S.s*. is found in transcriptomes measured by independent labs, this work suggests that intracellular Obelisk-*S.s*. populations are stably maintained in SK36. The evolutionary forces driving Obelisk persistence are SK36 remain unknown, and basic aspects of their evolutionary dynamics, such as Obelisk mutation rates, are not understood. It is unclear whether the Obelisk-*S.s*. polymorphisms reported here have any functional or adaptive significance, and the extent to which the dynamics of these polymorphisms are driven by selection is unknown.

We can speculate on possible mechanisms driving long-term Obelisk persistence in bacterial cells. One possibility is that Obelisks encode some beneficial function to the host cell. Alternatively, Obelisks could encode some post-segregational killing mechanism, much like the addictive toxin-antitoxin systems that stabilize many plasmids against stochastic loss. Such systems produce a short-lived antitoxin and a long-lived toxin; if the system is lost, the antitoxin is lost while the toxin persists, killing the cell^8^.

Our mathematical model suggests that high abundance can play a transient role in stabilizing Obelisk persistence in SK36, by reducing the stochastic loss rate of Obelisks from daughter cells. The model also shows that Obelisk persistence can be stabilized indefinitely when Obelisk abundance is so high that the probability of generating a daughter cell without any Obelisks goes to zero. Therefore, the simplest hypothesis for long-term Obelisk persistence, illustrated by our mathematical model, is that Obelisks may be so abundant in SK36 that they can persist in the absence of any benefit to its host.

Finally, another possibility is that Obelisk persistence is stabilized by horizontal gene transfer^9-12^. It is well established that *S. sanguinis* produces extracellular membrane vesicles^2,5^, and it has been reported that these vesicles contain small RNAs^2^. There is also some evidence that small RNAs can be transmitted between cells, although aspects of this hypothesis remain controversial^13,14^. Therefore, we can speculate that Obelisks may undergo intercellular transfer through some unknown mechanism, perhaps involving extracellular membrane vesicles.

Fundamentally, basic aspects of Obelisk biology and function are poorly understood. Understanding how this new class of circular RNA function and evolve remains an exciting problem at the frontier of the biological sciences.

## Methods

### Obelisk-S.s cDNA sequence

The following Obelisk-*S.s* cDNA sequence was reported by Zheludev et al. (2024) in Supplementary_table_1_stringent_Obelisk_clustering_011724.tsv, and used for downstream sequence analysis.

>Obelisk_000003|Obelisk-*S.s*.

ACTTGGTTAGTCCAGGAACTGTAATATATTAGAAAGGAAGTAAACCAAACATGTTAGATTGGAATACCTCATCAGACATCTTCGTCGAGAAGCTTCTTCAGAGAAACTACAAGAGTCAGAGTCTGCACAGCCAACCTCGCCATCGACCCCAAGTGGATGGAATTCCTTACGAGTTTGGATACAAAGGAACGATCTATCCTCTGAATAAATCACGAAACTGTATCATCATCTTGCTGTTGATACCCATTCTAGTTCACAGTACCAGAAATGCAGCCTACTTCGAGAGTCTCGAGAAGAAAATTGTCGAGCAAGTGAAGCTAAACAGGGCTCAAGGTAAATGGCAATTAGTCAGAGAACTTCTCGGACTAAAAGGCACTTTCCTCAAGCCCCGCTGGCAACACTTTGCGAAGACAGTTTCTTCAAGAGACTTCTTCGGAAATTGGCTACCTCTGATGCTAGAAATAGAAAGGTACCTTTACAGTAAAAAGATGTATCCAGATTCATATTTATCCTGGGACGATCATTCTTCGTACCGAGTTCGCAAGAAAGTCTACCGCCGTGGTTATGACGACAAGGGTAGCCGGAGACCTGAACACAAGTGGTTCCCTGAGAATGCCTTCTCTCGAGAATTGCTTGATGAAGTTCCGGTCAAACGTGCTGTTTACAAGCCGTTCGAACTATATCATGGTTACTCTGAAAAACGAAGGCGGAGATCGTCTCTAAGTTCTCTTCTAGATTTATAGGCAACGGAAAGCCTAAGAACTTAAGGTCGAACTTCTTCTTTCAAGAATTTCCTAATTGGTAAATTCTCTCAGTAAATCAATAACTTATTTTCCTTTGGAGAATTTGTTCCCTCTGAGGAAGAAGTAAAATTAAAATTTCGGGATTTTGAGGGACGGCAATCACGTCCCCTTTTCTCTTTTCGGAGAAAAAGGGATTGGGATTACCATCCCCTTAATTCTGTAAATTTTATTTTACCTCTTTCCTCTTAGGTTCAAATCCCCCAGAAGGGAAAAGTAGTTGACTATTGAAAGTCTCTACCCTATTGGGAAAGGCTCGTCGGAAAACAGTTCTCCGAAAGTTCTCATGACATTCTGGTGTCACAAAGTCGAGATGAGAACAAAGATTCGTCTTCGG

*RNAfold*^*15*^ 2.6.4 was used to predict the secondary structure of Obelisk-*S.s*.

### BLAST homology search

The NCBI BLAST web server^16^ (https://blast.ncbi.nlm.nih.gov/Blast.cgi?PROGRAM=blastn&PAGE_TYPE=BlastSearch&LINK_LOC=blasthome) was used to compare the Obelisk-S.s. cDNA sequence to the Nucleotide collection (nr/nt) database. The search was repeated, using the Organism field to restrict the homology search to *Streptococcus sanguinis* SK36 (taxid:388919).

### SK36 genome resequencing analysis

I manually annotated the location of the Oblin-1 in the Obelisk-*S.s* cDNA sequence and downloaded a Genbank-formatted reference sequence for Obelisk-*S.s* using Benchling^17^.

I downloaded a full Genbank-formatted reference sequence for the S. sanguinis SK36 genome from the NCBI RefSeq database^18^ at: https://www.ncbi.nlm.nih.gov/nuccore/NC_009009.1/

SK36 genome resequencing data was downloaded from the NCBI Short Read Archive (SRA) at: https://www.ncbi.nlm.nih.gov/sra/SRR14406732.

I then used *breseq*^19^ version 0.37.0 to jointly map the Illumina sequencing reads (<u>SRR14406732</u>) to the SK36 reference genome and the Obelisk-S.s. reference sequence, as follows:

breseq -j 10 -p -o ../results/SK36-DNAseq-breseq-polymorphism -r ../data/SK36-genome.gbk - r ../data/Obelisk_000003.gbk ../data/SRR14406732.fastq.gz

This was recorded in a shell script called *map-SK36-DNAseq-data.sh*.

### SK36 transcriptomic data and analysis

*pysradb*^20^ 2.1.0 and *SRA Toolkit* 3.0.5^21,22^ were used to fetch metadata and download transcriptomic data isolated from *S. sanguinis* SK36 monocultures (Tables 1 and 2).

*kallisto*^6^ was used to map the transcriptomic data for each sample in Table 1 to the SK36 reference genome and the Obelisk-S.s. reference sequence. This was automated with a Python 3.11 script called *run-kallisto-on-SK36.py*.

*breseq*^19^ version 0.37.0 was used to map the transcriptomic data for each sample in Table 1 to the SK36 reference genome and the Obelisk-S.s. reference sequence, using a shell script *map-SK36-RNAseq-data.sh. breseq* was run in polymorphism mode (-p option) with default parameter settings, which reports mutations at a 5% allele frequency threshold.

### Mathematical model

We built a mathematical model to examine how Obelisk copy number affects Obelisk evolutionary stability. A diagram of the model is shown in Figure 1B. This model involves two subpopulations of bacteria. The first carries an intracellular population of Obelisks (Type 1), and the second is an Obelisk-free subpopulation (Type 2). We are interested in the dynamics of these two populations due to selection (growth) and mutation (stochastic loss of intracellular Obelisks during cell division).

An interactive Pluto computational notebook of the model, called *obelisk-abundance-model.jl*, is available at: https://github.com/rohanmaddamsetti/obelisk-SK36/. This notebook can be run by installing and running Pluto.jl within Julia 1.10+ (see instructions at: https://plutojl.org/) and then opening the notebook using the Pluto web browser interface. Unless otherwise stated, the simulation results shown in Fig. 1 use the following default parameter settings (units of Obelisk copies per cell): Obelisk copy number *N* = 10, Obelisk capacity *N\** = 1000.

### Model assumptions

The population is modeled as a vector of two states, **x**, where *x*_*1*_ represents a subpopulation contain intracellular Obelisks, and *x*_*2*_ represents a subpopulation that has lost the Obelisks.

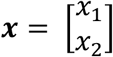

The total population size *X*(*t*) = *x*_*1*_(*t*) + *x*_*2*_(*t*).

### Selection dynamics

We assume that subpopulation *x*_*1*_ grows at rate *r*_*1*_, and subpopulation *x*_*2*_ grows at rate *r*_*2*_. We use the term “fitness” synonymously with these growth rates. We assume that *r*_*1*_ = *r*_*2*_ – α*N*, where *N* is the Obelisk copy number per cell. This assumption means that Obelisks have no fitness benefit and confer a small linear fitness cost α per copy. We define *N\** as the Obelisk capacity; that is, the Obelisk copy number at which *r*_*1*_ = 0 (Figure 1D). So,

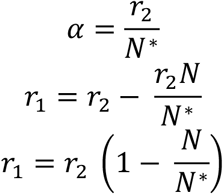

In our model, Obelisk copy number *N* and Obelisk capacity *N\** are free parameters, under the assumption that 0 ≤ *N* < *N\**.

### Mutation dynamics

We assume that *x*_*1*_ cells generate daughter cells at rate *r*_*1*_, and that those daughter cells fail to inherit any Obelisks with probability µ. We model µ as a geometrically decreasing function of Obelisk copy number *N*, based on the following argument:

Suppose a cell with Obelisks divides. The per capita rate of generating a cell without Obelisks is the probability that one daughter cell gets no Obelisks, and the other gets all of its parent’s Obelisks (Figure 1C). We assume partitioning is purely stochastic. So, the probability that a daughter doesn’t get an Obelisks is:

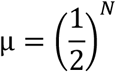

Since the per-capita growth rate for Type 1 cells is *r*_*1*_, the per-capita rate of generating Obelisk-free daughter cells is *r*_*1*_µ, and the per-capita rate of generating daughter cells with Obelisks is *r*_*1*_ (1 – µ).

### Full dynamics

The selection and mutation dynamics are combined into a matrix **A**:

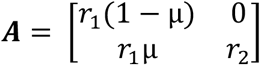

So, the full dynamics are modeled by the following matrix system of ODEs (Figure 1B):

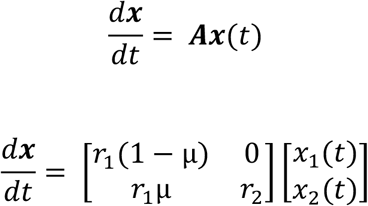

### Analytical solution for response time

Consider the initial condition 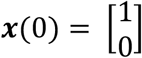 ; that is, *x*_*1*_(0) = 1 and *x*_*2*_(0) = 0. Since we assume that *r*_*1*_ *< r*_*2*_, eventually *x*_*1*_ ≤ *x*_*2*_. How long does it take for *x*_*2*_ = *x*_*1*_ ? This is a natural measure of the transient stability of the *x*_*1*_ state. We have a linear system of ODEs that can solved analytically:

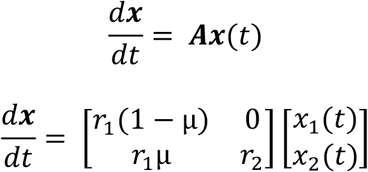

The eigenvalues of **A** are the diagonal entries *r*_*1*_(1–µ) and *r*_*2*_, as these are the roots of the characteristic polynomial of **A**. Solving for the corresponding eigenvectors, we find the following general solution:

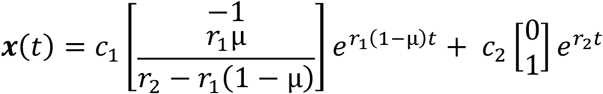

Using the initial condition 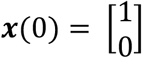, we find that:

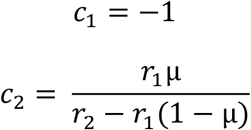

So the particular solution is:

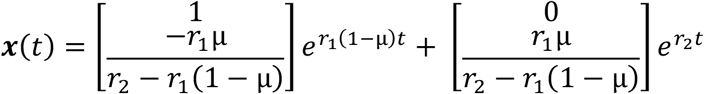

Now, we can solve for the response time *t\** where *x*_*1*_(*t\**) = *x*_*2*_(*t\**). Some algebra yields:

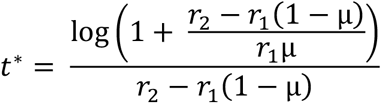

where log() is the natural logarithm (base *e*). Without loss of generality, we can let *r*_*2*_ = 1, so that 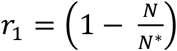 and 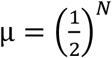 and examine how *t*\* changes as we change Obelisk copy number *N* and Obelisk capacity *N*\* (which defines the burden or fitness cost of an intracellular Obelisk population). This result is shown in Figure 1H.

### Data and Code Accessibility

All data analyzed in this work is publicly available. NCBI accessions are provided in the Tables and Methods. All data analysis code is available at: https://github.com/rohanmaddamsetti/obelisk-SK36.

## Acknowledgements

We thank Nkrumah Grant and Yasa Baig for discussions that motivated this work, and Jia Lu for helpful discussions about the mathematical model.

